# The receptor-like kinase ALE2 promotes giant cell formation in the sepal epidermis

**DOI:** 10.1101/2024.08.29.610297

**Authors:** Frances K. Clark, Jessica McGory, Nicholas J. Russell, Pau Formosa-Jordan, Adrienne H. K. Roeder

**Author notes:** Authors contributed equally to this work (Frances K. Clark, Jessica McGory). Author for correspondence: (AHKR).

## Abstract

During sepal development in *Arabidopsis thaliana*, epidermal pavement cells differentiate into small cells and large, highly endoreduplicated giant cells. While the placement of giant cells differs between sepals, the number of giant cells is fairly consistent. The HD-ZIP class IV transcription factor ATML1 has been found to promote giant cell formation. ATML1 protein fluctuates within epidermal nuclei of developing sepals and high ATML1 concentrations reached in the G2 phase of the cell cycle strongly correlates with giant cell fate specification. A genetic screen for reduced giant cell number identified the receptor-like kinase ABNORMAL LEAF SHAPE2 (ALE2) as being important for giant cell formation. We find that ALE2 functions genetically upstream of ATML1 in promoting the formation of giant cells. We observe that nuclear-localized mCitrine-ATML1 fluctuates in *ale2* mutants much as it does in wild type and, importantly, nuclear mCitrine-ATML1 reaches similarly high peak concentrations in *ale2-1* mutants as in wild type. This indicates that ALE2 functions upstream of ATML1 by affecting protein activity rather than gene expression or protein degradation. One function of ATML1 is to promote transcription of the CDK inhibitor LGO. We find that LGO transcription is delayed and decreased in *ale2-1* mutants as compared to wild-type plants, consistent with ATML1 having impaired protein function in *ale2* mutants. Overall, we find that the receptor-like kinase ALE2 is necessary to sensitize cells of the developing sepal epidermis to fluctuations of the transcription factor ATML1.

## Introduction

How complex patterns of cell types form from a small number of identical progenitor cells is a fundamental question in developmental biology. Patterning in nature often relies on a combination of cell-cell signaling and stochasticity. For example, in *Drosophila* ommatidia differentiation, random variation in expression levels of the transcription factor Spineless drive R7 photoreceptor cells to take on a “pale” identity (low expression) or a “yellow” identity (high expression) (Wernet et al., 2006). The specified “yellow” R7 photoreceptor cells then impose the “yellow” identity on R8 photoreceptor cells whose default identity is “pale” (Wernet et al., 2006). The result is that R7 and R8 photoreceptor cells within the same ommatidia have the same identity and the “pale” and “yellow” ommatidia are scattered randomly across the compound eye. In plants, trichomes are patterned such that they do not touch one another and are regularly spaced across the leaf surface (Hulskamp, 2004). The mechanism underlying this patterning is consistent with a combined reaction-diffusion and activator depletion model that relies on initial stochastic differences in an activator complex between identical cells (Balkunde et al., 2020; Hulskamp, 2004). The activator complex promotes the expression of mobile inhibitors that pass into surrounding cells to inhibit trichome formation in a form of intercellular communication (Digiuni et al., 2008).

The *Arabidopsis thaliana* (henceforth Arabidopsis) sepal has an outer epidermis composed of cells of different sizes. Specialized cells like stomatal guard cells and trichomes have characteristic sizes, with the former being small and the latter being large. Interestingly, the other epidermal cells that serve to provide a barrier between the environment and internal plant cell layers, called pavement cells, have a wide range of sizes. On the sepal epidermis the largest of these pavement cells are called giant cells (Roeder et al., 2010). Giant cells are not only very large but are highly endoreduplicated. Endoreduplication occurs when cells circumvent the traditional G1-S-G2-M cell cycle and instead enter a cycle of DNA replication and growth, bypassing the division process. Giant cells endoreduplicate multiple times and are typically 16C and above (Roeder et al., 2010; Roeder et al., 2012). In addition to their large size and high ploidy level, giant pavement cells are distinct from smaller pavement cells in that their transcriptome indicates a function in insect and pathogen defense (Schwarz & Roeder, 2016). Giant cell localization varies from sepal to sepal and initiates randomly with giant cells touching one another sometimes (Clark et al., 2024). Importantly, the number of giant cells in wild-type sepals falls within a narrow range, which appears to be necessary for proper sepal curvature (Roeder et al., 2012). How giant cells are patterned during development such that their location varies but their number is fairly conserved is an essential question.

Previous research in our lab has shown that fluctuations of the HD-ZIP Class IV transcription factor ATML1 play a role in giant cell patterning (Meyer et al., 2017). ATML1 protein concentration fluctuates in the nuclei of early-developing sepals. High concentrations of ATML1 reached during the G2 phase of the cell cycle correlates strongly with giant cell differentiation. This data led to a model in which ATML1 fluctuates randomly and ATML1 surpassing a concentration threshold in the G2 phase leads to giant cell differentiation. This model was supported by simulations showing that it could recapitulate several dynamical aspects of the giant cell fate commitment process, and it could also produce a scattered pattern of giant cells (Meyer et al., 2017). If the probability of ATML1 surpassing the concentration threshold was consistent, then wild-type sepals would be expected to have similar numbers of giant cells. If the ATML1 fluctuations were stochastic, then giant cell localization would be expected to vary from sepal to sepal in a seemingly random manner. By comparing time-lapse data of developing sepals and the model for giant cell fate decisions, we have recently shown that giant cells appear randomly throughout the epidermis, and cell divisions cause giant cells to become more clustered than expected by chance during development (Clark et al., 2024).

Although this model explains a lot about giant cell patterning, there are additional genes that we isolated through a forward genetics screen involved in giant cell specification. In addition to ATML1, the receptor-like kinase ACR4, the calpain protease DEK1, and the cyclin-dependent kinase (CDK) inhibitor LGO were found to promote giant cell formation (Roeder et al., 2010; Roeder et al., 2012). Double mutant analysis shows that these genes are in the same genetic pathway and that ACR4 is genetically upstream of ATML1, whereas DEK1 and LGO act genetically downstream of ATML1 (Meyer et al., 2017). The fact that a receptor-like kinase is involved in giant cell differentiation opens up the possibility that cell-cell signaling is involved in giant cell patterning.

In addition to their role in giant cell specification, ACR4, ATML1, and DEK1 are involved in epidermal specification (Abe et al., 2003; Gifford, Dean, & Ingram, 2003; Johnson et al., 2005; Watanabe et al., 2004). Without epidermal specification, the Arabidopsis embryo cannot progress past the globular stage of development (Johnson et al., 2005; Ogawa et al., 2015). Like many other developmental regulators, these genes play more than one role. After specifying epidermal identity, they pattern giant cell formation in sepals.

Here we report that the receptor-like kinase ABNORMAL LEAF SHAPE2 (ALE2) is also involved in giant cell patterning. Just as ACR4, ATML1, and DEK1, ALE2 plays a role in epidermal specification (Tanaka et al., 2007). The *ale2-1* loss-of-function allele is caused by a single base pair insertion that creates a frameshift disrupting the kinase domain (Tanaka et al., 2007). Homozygous *ale2-1* mutants have epidermal defects, including disorganization of epidermal tissues, defects in the leaf cuticle, and fusion of floral organs (Tanaka et al., 2007). *ale2-1* mutants are also sterile due to malformation of the ovules and decreased pollen formation (Tanaka et al., 2007). We find that *ale2* mutants also lack sepal giant cells. Here, we investigated how a signaling pathway fits into giant cell specification by investigating how this receptor-like kinase functions to promote giant cell formation.

## Results

### *ALE2* is required for giant cell formation

We performed a forward genetic screen for mutants affecting the development of giant cells and identified the *ale2-3* mutant in an ethyl methanesulfonate (EMS) mutagenized Landsberg *erecta* (L*er)* M2 population (Figure 1A–F) (Roeder et al., 2010; Roeder et al., 2012). *ale2-3* exhibited a complete loss of giant cells (Figure 1E,F). The *ale2-3* mutation was positionally cloned and identified as disrupting the gene encoding the transmembrane receptor-like serine/threonine-protein kinase ALE2. Our novel *ale2-3* mutation is the result of a G-to-A change at position 849 of the coding sequence resulting in a nonsense mutation that introduces a premature stop codon at amino acid 283, just after the transmembrane domain (Figure 1M). We found that giant cells are also lost in the previously published *ale2-1* allele in the Columbia-0 (Col-0) wild-type background (Figure 1G–L) (Tanaka et al., 2007). We used the *ale2-1* allele for the remainder of our work. The *ale2-1* allele is the result of a frameshift mutation due to a single nucleotide insertion in the seventh exon and at amino acid position 283 (Figure 1M). We were able to rescue the *ale2-1* mutant phenotype by expressing wild-type *ALE2* under the ubiquitous 35S promoter, confirming that the defects observed were the result of the *ale2* mutation (Supplemental Figure 1A–C).

**Figure 1:**
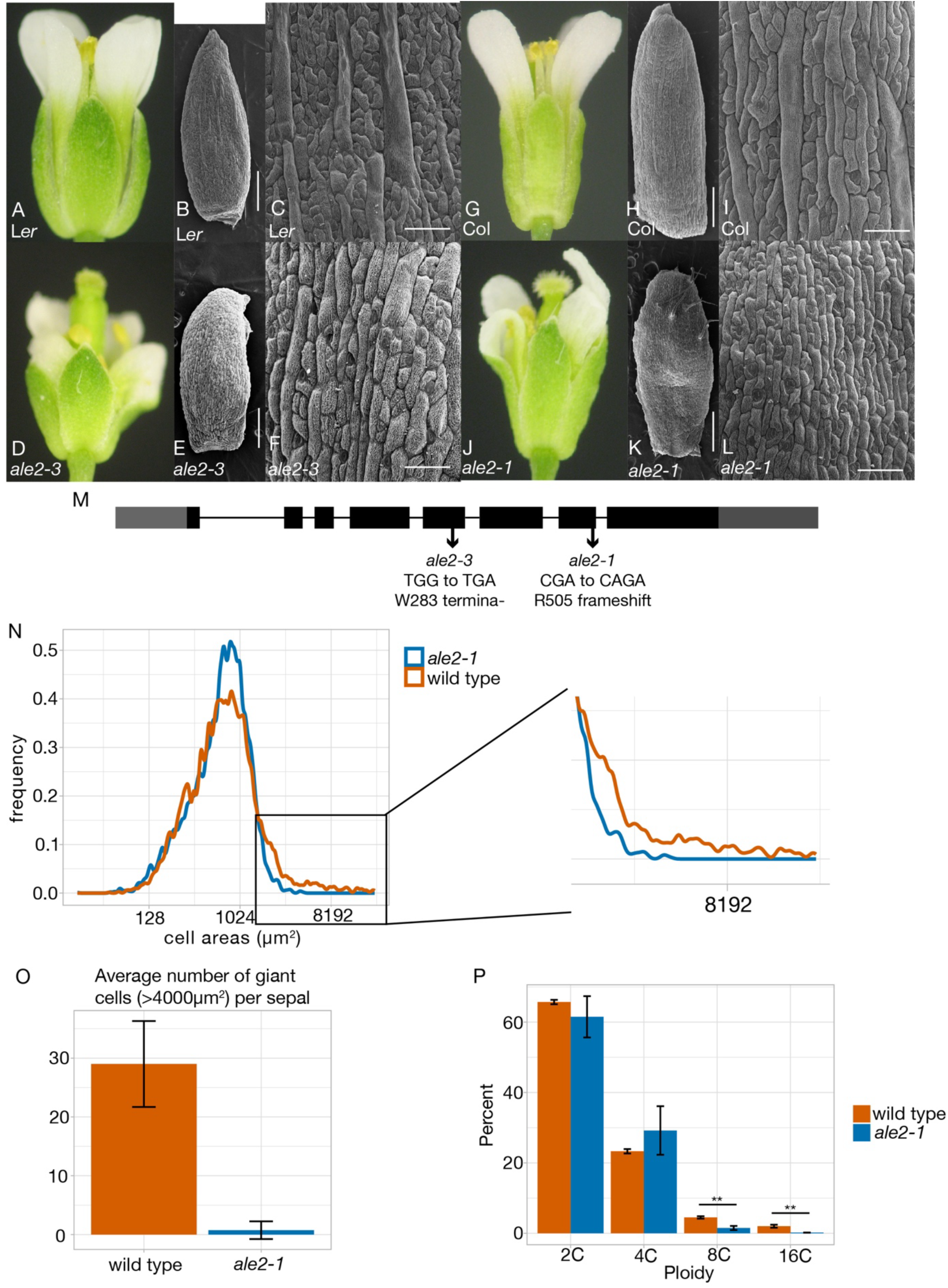
*ALE2* is necessary for sepal giant cell formation. (A) Columbia-0 (Col-0) wild-type flower. (B–C) SEM images of Col-0 wild-type sepal. Scale bar for B is 250 µm and for C is 50 µm. (D) *ale2-1* flower in Col-0 background. (E-F) SEM images of *ale2-1* sepal in Col-0 background. Scale bar for E is 250 µm and for F is 50 µm. (G) Landsberg erecta (L*er)* wild-type flower. (H-I) SEM images of wild-type sepal in L*er* background. Scale bar for H is 250 µm and for I is 50 µm. (J) *ale2-3* flower in L*er* background. (K-L) SEM images of *ale2-3* sepal in L*er* background. Scale bar for K is 250 µm and for L is 50 µm. (M) *ALE2* gene map including locations of *ale2-1* and *ale2-3* mutations. (N) Cell areas of wild-type and *ale2-1* mutant sepals. Density plot includes abaxial epidermal cells from 4 *ale2-1* sepals and 5 wild-type sepals. (O) Average number of giant cells per wild-type Col-0 and *ale2-1* sepals. Graph made using the same data as in N. Wild type has a higher average number of giant cells according to a 2 sample t-test assuming equal variance (t(7)=-7.50, p= 0.00014). (P) Percentage of cell ploidy in wild-type and *ale2-1* sepals. Epidermal cells from both the abaxial and adaxial side of sepals included. Wild type has a higher percent of 8C cells according to a 2 sample t-test assuming unequal variance (t(3) =6.73, p=0.0067) as well as a higher percent of 16C cells according to a 2 sample t-test assuming unequal variance (t(3) =7.39, p=0.005).

To examine the epidermal pattern of these mutant sepals more closely, we quantified epidermal cell areas in confocal projection images of the mature sepals. In *ale2-1*, the cell size distribution is altered such that the largest cells, the giant cells, are absent, and the frequency of smaller cells increases (Figure 1N). Cell area scales with ploidy (Jorgensen & Tyers, 2004; Melaragno, Mehrotra, & Coleman, 1993) and a giant cell can be defined as having an area larger than 4,000 µm^2^ (Meyer et al., 2017; Roeder et al., 2012). Under this definition, wild-type sepals have an average of 29 giant cells per sepal, whereas *ale2-1* mutants have an average of less than one giant cell per sepal (Figure 1O). The largest recorded cell area in *ale2-1* was 4,096 µm^2^ compared to 22,122 µm^2^ in wild type. A typical giant cell has a ploidy of about 16C. We used flow cytometry to measure ploidy in the sepal epidermis and found 0% of cells were 16C in *ale2-1* mutant sepals, compared to 2% of cells in wild-type sepals (Figure 1P). Thus, we confirm that giant cells are absent in *ale2-1* mutants.

We also wondered whether ALE2 affects the size of cells outside of the sepal. Genes affecting sepal giant cell formation have also been found to affect cell size patterning in the leaf epidermis (Clark et al., 2024). We imaged leaf 1 or 2 25 days post germination (Leaves 1 and 2 initiate at the same time and are phenotypically equivalent). Images were taken on the abaxial face of the leaf midway between the midrib and margin and midway between the tip and base and cell areas were quantified. Wild-type leaves have more larger cells than *ale2-1* leaves (Figure 2 A-G). The average number of cells above a leaf giant cell area threshold of 14,160 µm^2^ (Clark et al., 2024) established was significantly greater for wild type (mean=15, n=3) than for *ale2-1* (mean=4, n=3; two-sample 2-tailed t-test t(3.7)=3.81, p=0.022).

**Figure 2:**
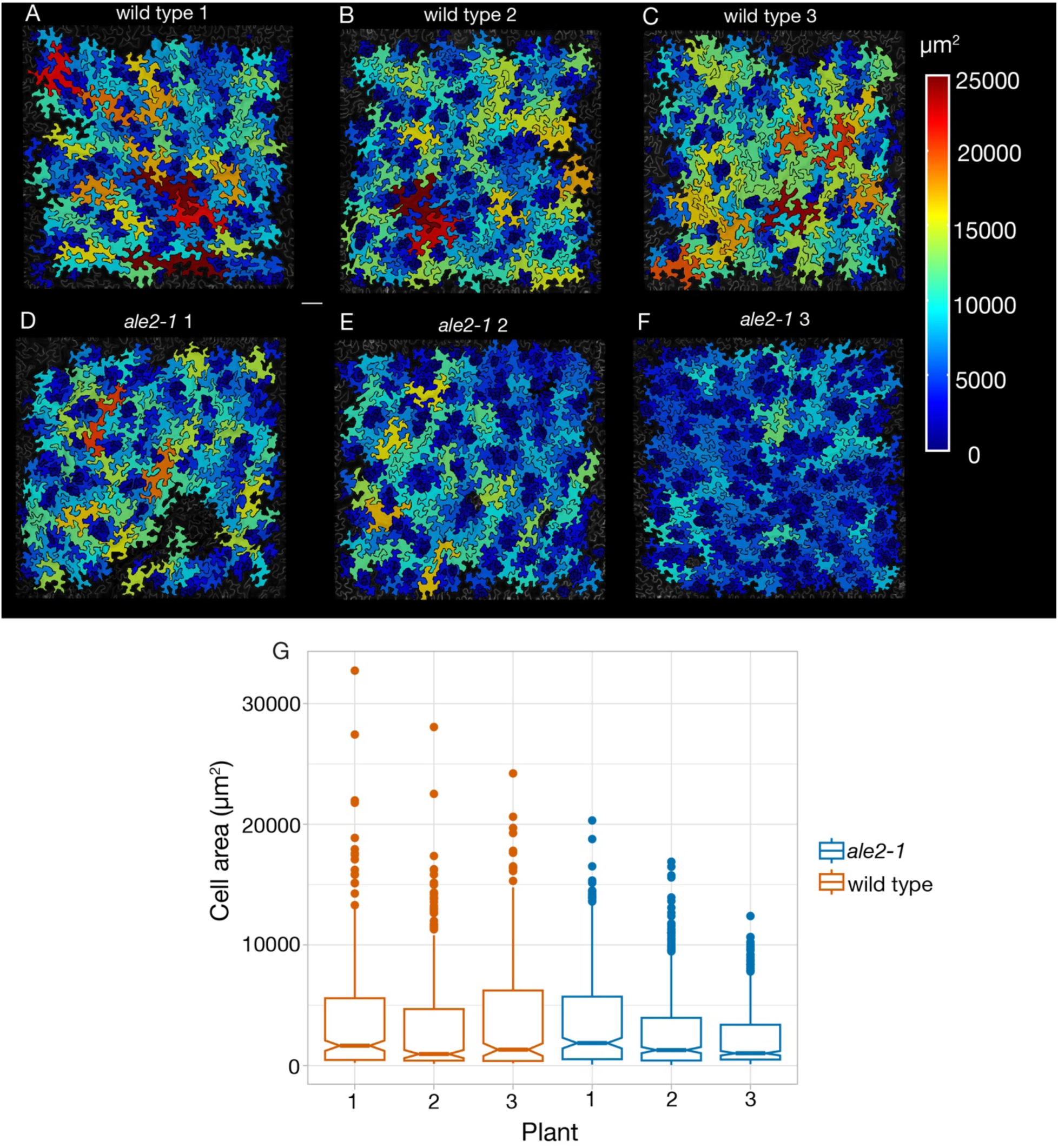
ALE2 promotes giant cell development in the leaf. Cell area heat maps in µm^2^ of leaf 1 or 2 at 25 days post germination for (A–C) wild type and (D–F) *ale2-1* mutant. Scale bar is 100 µm. (G) Cell areas of wild-type and *ale2-1* replicates plotted as boxplots. Replicate in A used in (Clark et al., 2024). Replicates in A, B, and C used in (Trozzi et al., 2023).

Previously, we have demonstrated that giant cell identity, in terms of specific patterns of gene expression, can be separated from giant cell size (Roeder et al., 2012). Because our data shows that *ale2-1* mutants fail to differentiate enlarged polypoid giant cells, we wondered whether giant cell identity is also lost. We examined the expression of two fluorescent markers corresponding to a giant cell enhancer reporter and a small cell enhancer reporter (Hong et al., 2023; Roeder et al., 2012) in wild type and *ale2-1*. The *ale2-1* mutant produced mature sepals with very few cells that express the giant cell enhancer reporter and more cells expressing the small cell enhancer reporter (Figure 3A, B). This phenotype is similar to the *dek1-4* mutant, which also exhibits a significant reduction in giant cell enhancer reporter expression along with loss of enlarged cells (Roeder, 2012). However, it is distinct from the *lgo* mutant phenotype, in which, despite the loss of giant cells, the giant cell enhancer reporter is still prominently expressed in several small cells (Roeder et al., 2012). Even so, in earlier stages of sepal development, *ale2-1* mutants often have a small number of medium and small cells containing the giant cell enhancer reporter signal (Figure 3A1–A2, B1–B2). It is possible that these few cells attempt to differentiate as giant cells early on, and thus express the giant cell enhancer reporter, but are unsuccessful in triggering the endoreduplication pathway and therefore never become phenotypically enlarged polyploid giant cells. Notably, some slightly enlarged cells in *ale2-1* expressed the giant cell enhancer marker later in development (Figure 3B3).

**Figure 3:**
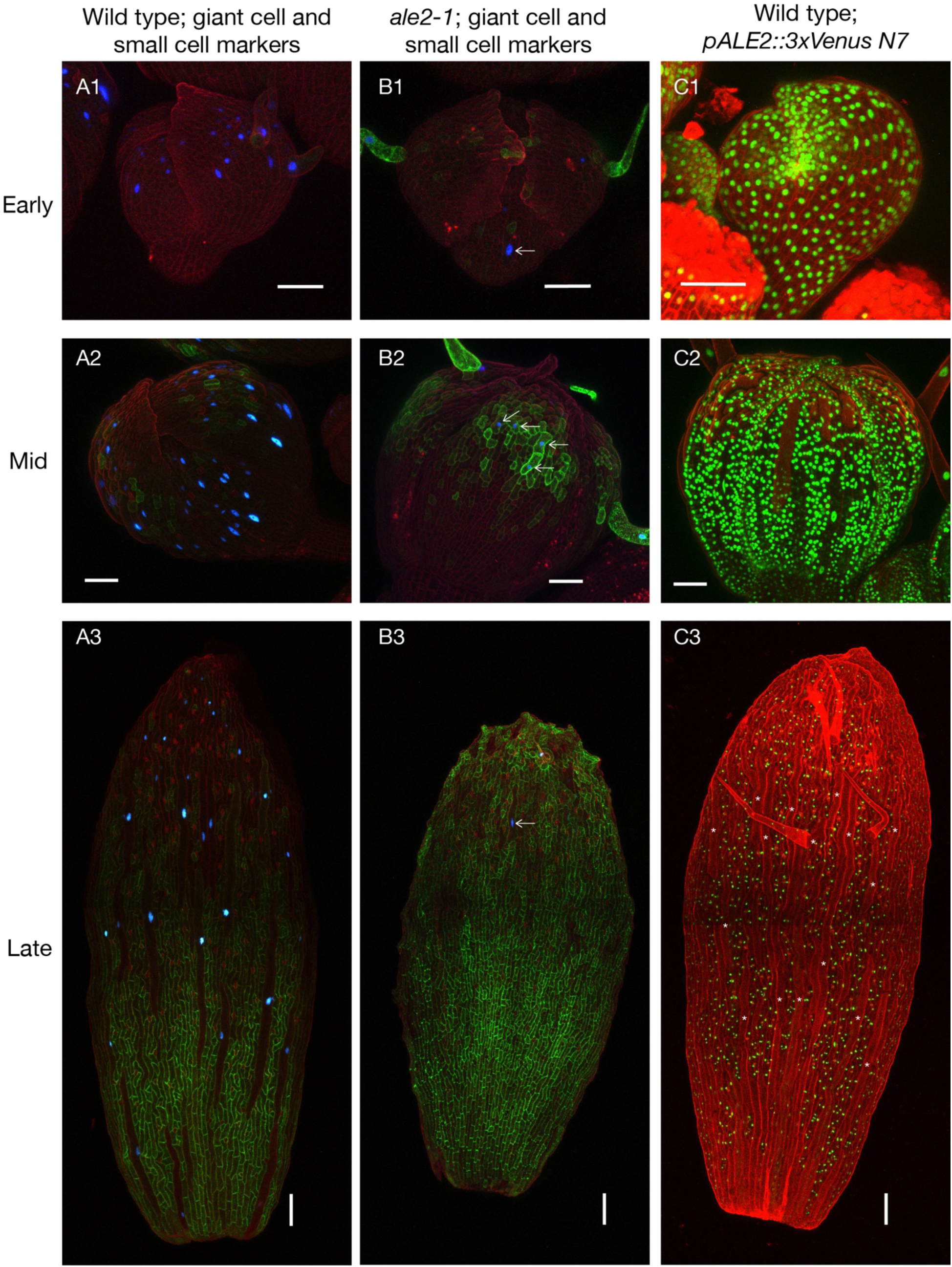
*ale2-1* sepals lose giant cell identity. (A1–A3) Giant cell and small cell fluorescent markers in wild-type sepals at early, mid, and late stage development. Cell walls stained with propidium iodide (shown in red). Giant cell marker is nuclear localized 3XVenus-N7 (shown in blue) under the control of a giant cell specific promoter (Roeder et al., 2012). Small cell marker is ER localized GFP (shown in green) under the control of a small cell enhancer (Roeder et al., 2012). (B1–B3) Giant cell and small cell fluorescent markers in *ale2-1* sepals at early, mid, and late stage of development. Arrows indicate cells with giant cell identity in *ale2-1* mutant sepals. (C1-C3) ALE2 promoter driven expression in early, mid, and late stage of development (*pALE2:: 3xVenus-N7*, shown in green). Asterisks in C3 indicate giant cells. Early stage buds were at stages 6 or 7. Mid stage buds were at stages 9 to 10. Scale bars for A1, B1, C1, A2, B2 and C2 are 50 µm. Scale bars for A3, B3 and C3 are 100 µm.

### *ALE2* is expressed in all epidermal cells of developing sepals but its expression declines in giant cells after specification

Since *ale2* mutants were unable to form giant cells, we hypothesized that *ALE2* would be expressed in the progenitors of giant cells. Indeed, we saw that an *ALE2* promoter driving VENUS expression (*pALE2::3xVENUS-N7*) was active in all epidermal cells of the developing sepal before and during giant cell specification (Figure 3C1). *ALE2* promoter activity continued in newly differentiated giant cells but was subsequently lost (Figure 3C2). By Stage 12, *ALE2* promoter activity is absent in all giant cells (identified by enlarged nuclei and cell size), while smaller cells continue to have *ALE2* promoter activity (Figure 3C3).

We wondered whether the loss of *ALE2* expression in newly differentiated giant cells is necessary for maintenance of giant cell identity and differentiation. We tested this possibility by forcing *ALE2* expression after giant cell differentiation by putting it under control of the giant cell enhancer. When the giant cell enhancer drives expression of ALE2, giant cell number and patterning appear normal (Supplemental Figure 1D,E). Thus, ALE2 downregulation does not appear to be necessary for maintaining giant cell patterning.

### *ALE2* functions genetically upstream of *ATML1* and *LGO* to promote giant cell formation

To determine whether *ALE2* functions genetically in the giant cell differentiation pathway, we performed a double mutant analysis (Figure 4). Both *ale2-1 lgo2-1* and *ale2-1 acr4-2* double mutant sepals displayed a phenotype similar to *ale2-1* single mutants with a complete loss of giant cells (Figure 4B, F, J). These mutants also exhibited epidermal defects in organ structure similar in severity to the *ale2-1* single mutant. Based on these results, we conclude that ALE2 functions in the same pathway as ACR4 and LGO. To determine where ALE2 functions within this pathway, we crossed *ale2-1* mutants with plants overexpressing *ATML1* or *LGO*. In previous studies, we have shown that overexpressing *ATML1* (*ATML1-OX*) and overexpressing *LGO* (*LGO-OX*) results a huge increase in giant cells (Meyer et al., 2017; Roeder et al., 2012; Schwarz & Roeder, 2016). *ale2-1 ATML1-OX* sepals looked the same as *ATML1-OX* sepals in terms of still having many ectopic giant cells (Figure 4G, H). Likewise, *ale2-1 LGO-OX* sepals looked the same as *LGO-OX* sepals in terms of ectopic giant cells (Figure 4C, D). Thus, we conclude that *ALE2* is genetically upstream of both *ATML1* and *LGO*. From previous work, we additionally know that *LGO* acts downstream of *ATML1* (Schwarz, 2016), and with this information, we were able to form a pathway for giant formation that includes *ALE2* (Figure 4K).

**Figure 4:**
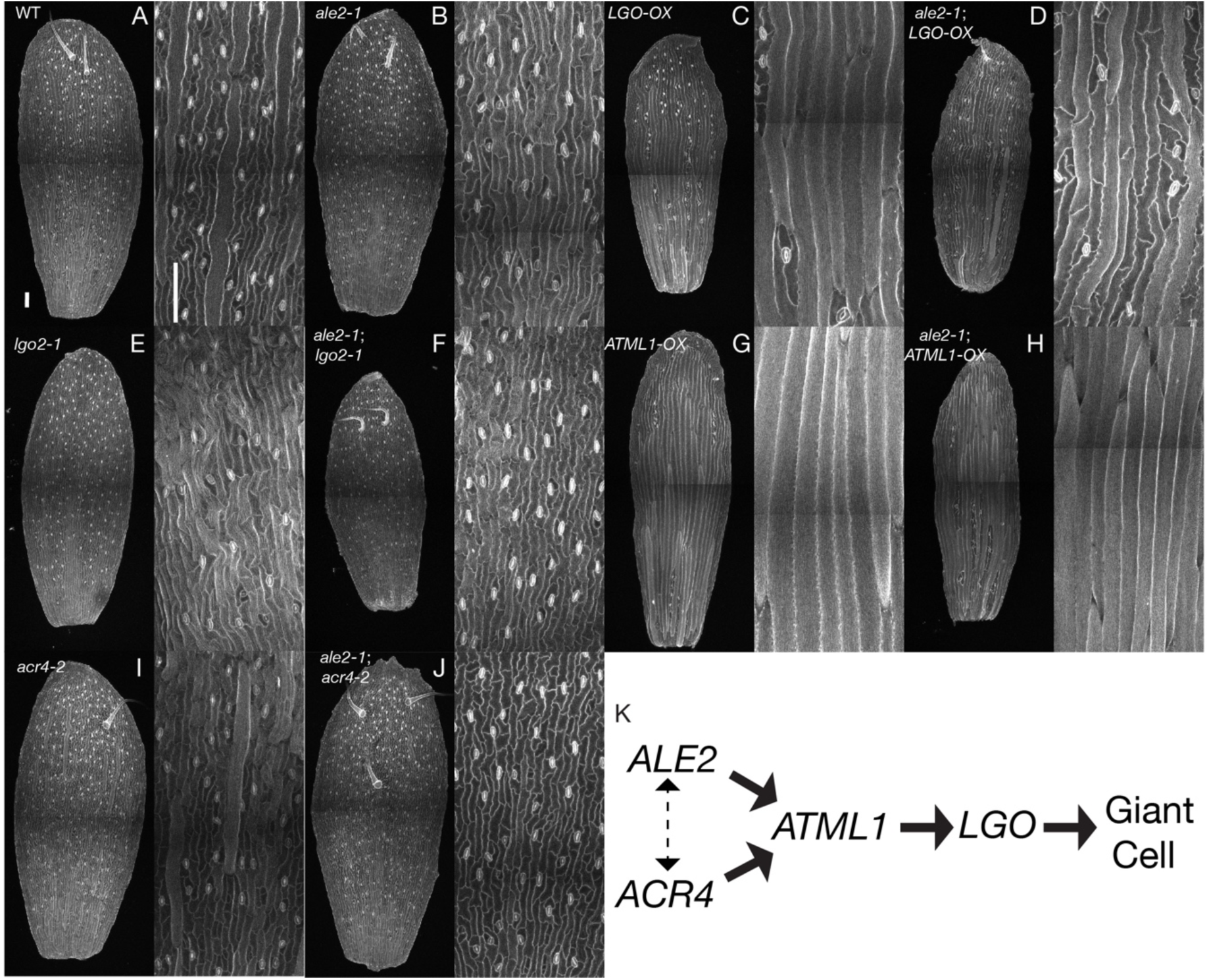
Double mutant analysis places *ALE2* upstream of *ATML1* and *LGO* in giant cell specification. (A–J) Abaxial sides of stage 12 sepals with cell walls stained with propidium iodide. (A) Wild type, (B) *ale2-1*, (C) *LGO-OX*, (D) *ale2-1 LGO-OX*, (E) *lgo-2*, (F) *ale2-1 lgo-2*, (G) *ATML1-OX*, (H) *ale2-1 ATML1-OX*, (I) *acr4-2* and (J) *ale2-1 acr4-2*. (a-j) Magnified images of the abaxial side of stage 12 sepals. (a) Wild type, (b) *ale2-1*, (c) *LGO-OX*, (d) *ale2-1 LGO-OX*, (e) *lgo-2*, (f) *ale2-1 lgo-2*, (g) *ATML1-OX*, (h) *ale2-1 ATML1-OX*, (i) *acr4-2* and (j) *ale2-1 acr4-2*. (K) Location of ALE2 in the giant cell formation pathway based on double mutant analysis. Normal arrows indicate genetic activation, whereas dashed arrows indicate potential interplay.

We were unable to generate double mutants between the loss of function *ale2-1* mutant and loss of function *atml1-3* mutant. Because *ALE2* and *ATML1* are both also involved in epidermal specification, we hypothesized that double mutant embryos might arrest at the globular stage of development, as other mutant embryos with severe epidermal specification defects have been found to do (Johnson et al., 2005; Ogawa et al., 2015; San-Bento et al., 2014; Tanaka et al., 2007). We examined developing embryos within the self-fertilized siliques of *ale2-1/+;atml1-3/atml1-3* plants to look for embryonic arrest (Figure 5). We found that in siliques from *ale2-1/+;atml1-3/atml1-3*, 76.1% of embryos were at the torpedo or walking stick stages (Figure 5D) while 23.9% of embryos were arrested at the globular stage (Figure 5C,E) (n=15 siliques), consistent with the expected 25% segregation of double mutant embryos (two-sample t-test: 2-tailed, df=24, t=-0.93764, p=0.3578, not significantly different from 25% expected by Mendelian genetics). In many of the arrested embryos, we observed that the epidermal layer was not smooth, which has previously been found in mutant embryos with a loss of epidermal identity (Johnson et al., 2005). As a control, we found that only 1.0% of *atml1-3* single mutant embryos were arrested (n=11 siliques). Thus, we conclude that *ale2-1;atml1-3* double mutants arrest at the globular stage.

**Figure 5:**
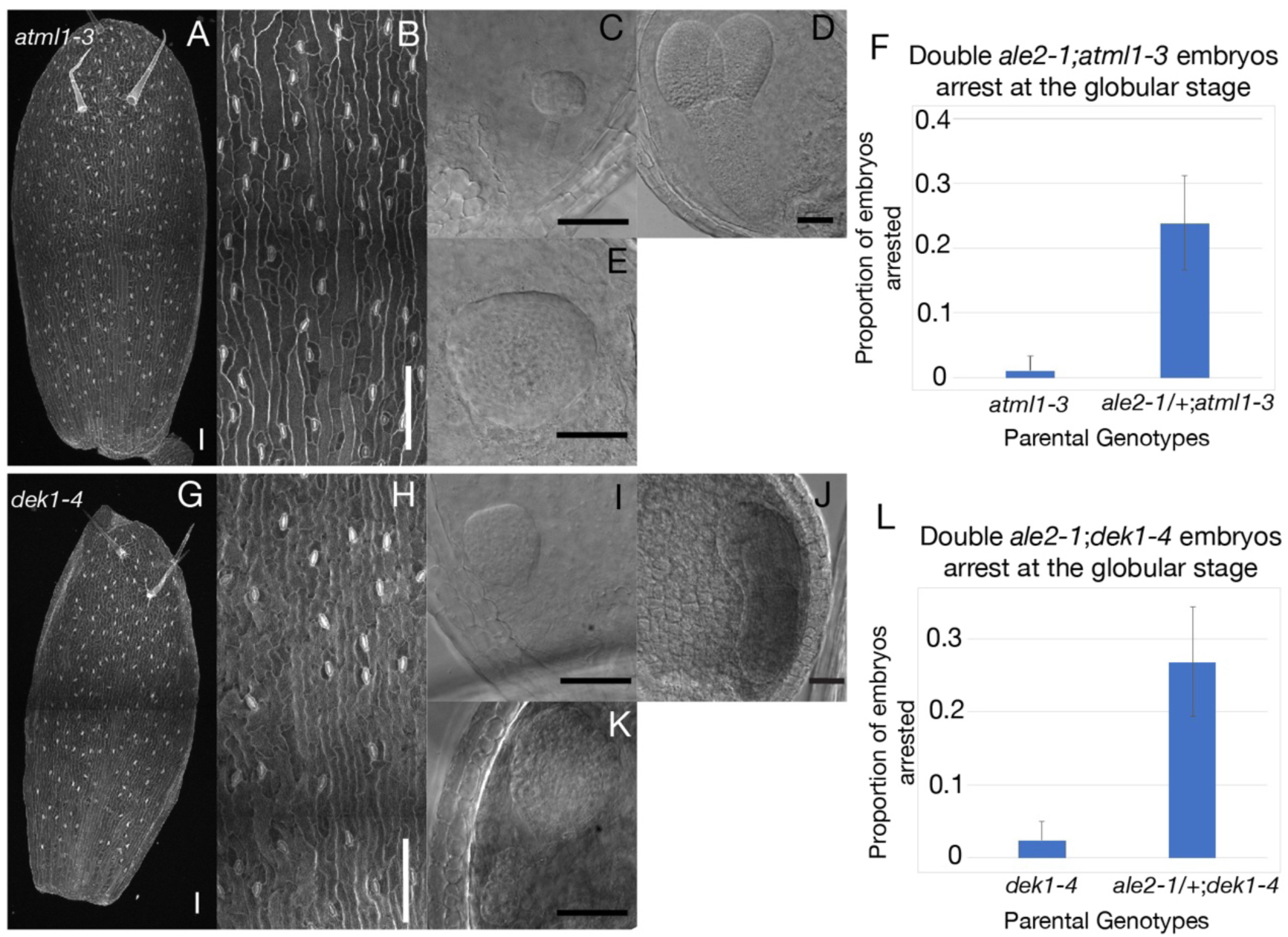
*ale2 atml1* and *ale2 dek1* double mutants exhibit embryonic arrest. (A) *dek1-4* mutant sepal PI stained, abaxial side. Scale bar=100 μm. (B) *dek1-4* mutant magnified sepal cells PI stained, abaxial side. Scale bar=100 μm. (C) Embryo arrested at the globular stage from a silique of a *dek1-4/dek1-4;ale2-1/+* plant. Scale bar=50 μm. (D) Normally developing embryo at the torpedo stage from a silique of a *dek1-4/dek1-4;ale2-1/+* plant. Scale bar=50 μm. (E) Embryo arrested and exhibiting abnormal growth from a silique of *dek1-4/dek1-4;ale2-1/+* plant. Scale bar=50 μm. (F) About 25% of embryos arrest in *dek1-4/dek1-4;ale2-1/+* plants. (G) *atml1-3* mutant sepal PI stained, abaxial side. Scale bar=100 μm. (H) *atml1-3* mutant magnified sepal cells PI stained, abaxial side. cale bar=100 μm. (I) Embryo arrested at the globular stage from a silique of an *atml1-3/atml1-3;ale2-1/+* plant. Scale bar=50 μm. (J) Normally developing embryo at the torpedo stage from a silique of an *atml1-3/atml1-3;ale2-1/+* plant. Scale bar=50 μm. (K) Embryo arrested and exhibiting abnormal growth from a silique of an *atml1-3/atml1-3;ale2-1/+* plant. Scale bar=50 μm. (L) About 25% of embryos arrest in *atml1-3/atml1-3;ale2-1/+* plants.

We were also unable to generate double mutants between *ale2-1* and the hypomorphic allele *dek1-4*. Since *DEK1* is also involved in epidermal specification, we examined the contents of *ale2-1/+;dek1-4/dek1-4* siliques to see if there was embryonic arrest at the globular stage as well. 26.9% of embryos from *ale2-1/+;dek1-4/dek1-4* siliques arrested at the globular stage (Figure 5I,K) (n=9 siliques), which was not significantly different from the expected 25% double mutant embryos (two-sample t-test: 2-tailed, df=12, t=-0.16226, p=0.8738, not significantly different from 25%), even after taking into account that 2.5% of *dek1-4* single mutant control embryos arrested (n= 5 siliques). Interestingly, a small number of both *ale2-1;atml1-3* and *ale2-1;dek1-4* embryos exhibited abnormal growth before arresting (Figure 5E,K). These embryos grew more than a typical globular embryo but were not able to initiate cotyledon formation. Such *ale2-1;atml1-3* embryos appeared spherical (Figure 5E) and such *ale2-1;dek1-4* embryos were typically oblong and had abnormal suspensor divisions (Figure 5K).

The epidermal lethal phenotype of *ale2-1;atml1-3* mutants, which is more severe than the epidermal defects of *ale2-1* and *atml1-3* single mutants, may point to *ALE2* being upstream of another gene besides *ATML1* important for epidermal specification. We speculate that *ALE2* might be upstream of the *ATML1* paralog *PDF2* as well. Double *atml1;pdf2* loss of function mutants are embryonic lethal and arrest at the globular stage (San-Bento et al., 2014) (Abe et al., 2003) (Ogawa et al., 2015). Arrest of *ale2-1;dek1-4* mutants indicates that loss of function of *ALE2* worsens the epidermal specification defect of hypomorphic *dek1* mutants. Loss of function *dek1* mutants are also embryonic lethal and arrest at the globular stage (Johnson et al., 2005).

### ALE2 does not affect ATML1 protein levels

Since ATML1 is downstream of ALE2, we wondered how ALE2 affects ATML1 to promote giant cell differentiation. We have previously shown that ATML1 nuclear protein concentrations fluctuate within individual nuclei of the developing sepal epidermis (Meyer et al., 2017). In addition, we have shown that high nuclear ATML1 protein concentration during G2 is strongly correlated with giant cell differentiation (Meyer et al., 2017). We initially hypothesized that ALE2 would be necessary for ATML1 to reach peak concentrations high enough to trigger giant cell differentiation, either by promoting *ATML1* expression or stabilizing ATML1 protein. To test this, we looked at mCitrine-ATML1 concentration in young buds directly preceding giant cell differentiation in wild type (*pATML1::mCitrine-ATML1; atml1-3,* in which the *atml1* mutation is rescued by the transgene) versus *ale2-1* mutant (*pATML1::mCitrine-ATML1; atml1-3; ale2-1*) (Figure 6A). Wild-type and *ale2-1* buds at Stage 4 both showed a range of nuclear mCitrine-ATML1 concentrations with similar distributions (Figure 6B). Critically, both wild type and *ale2-1* mutant nuclei showed similarly high peak concentrations of nuclear mCitrine-ATML1 (Figure 6B). This result contradicts our hypothesis and instead suggests that ALE2 does not regulate ATML1 concentration.

**Figure 6:**
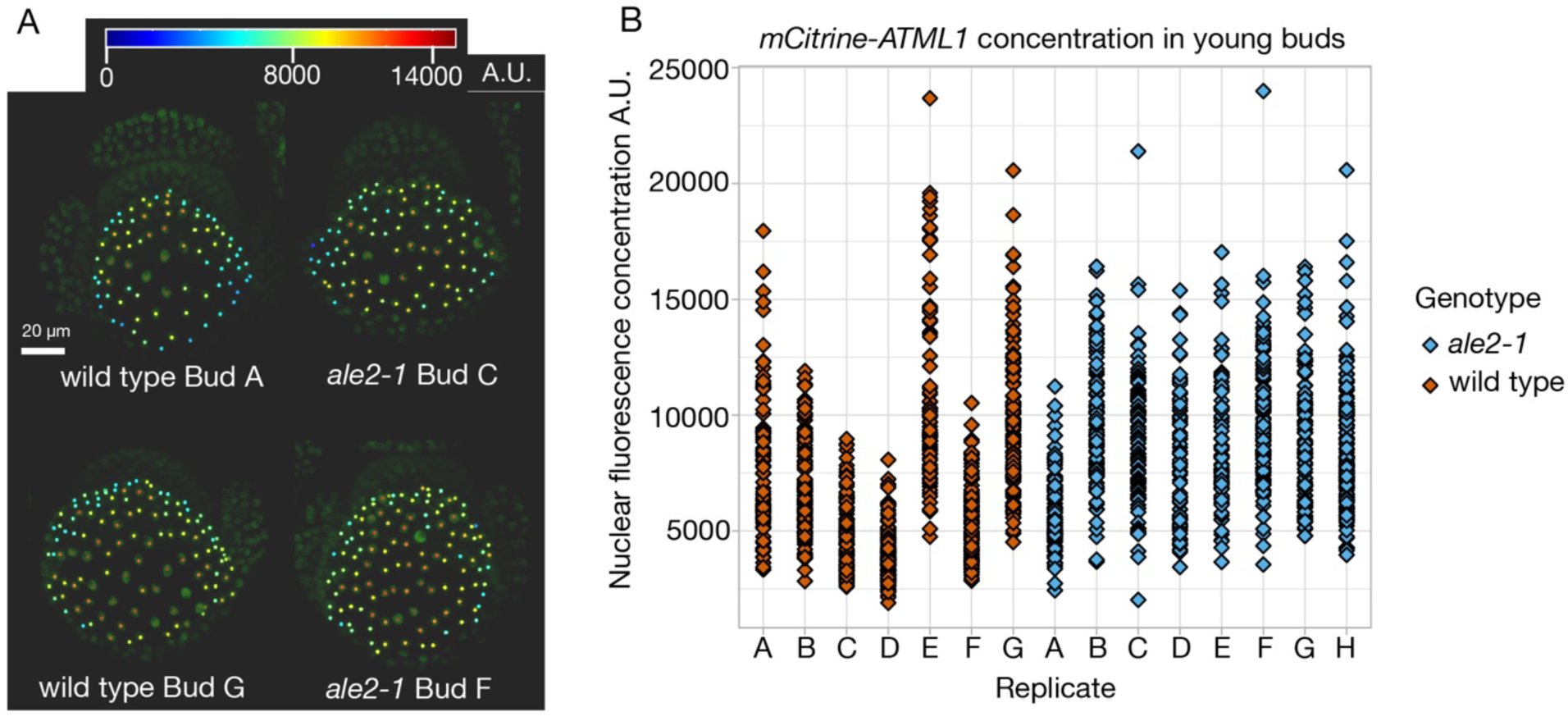
ALE2 does not affect ATML1 concentration peaks or fluctuations. (A) mCitrine-ATML1 protein concentration in 7 wild-type and 8 *ale2-1* mutant sepals on young flower buds (*pATML1::mCitrine-ATML1* atml1-3 background). Each dot represents a single nuclear ATML1 concentration. (B) Heat maps of mCitrine-ATML1 protein concentration for two representative wild-type buds (left) and two representative *ale2-1* mutant buds (right). Bud letter IDs can be referenced to the graph in A. Scale bar is 20 µm.

### ALE2 is important for proper *LGO* expression at the critical time of giant cell differentiation

Because peak ATML1 protein concentration is unaffected in *ale2-1*, we wondered whether ALE2 works upstream of ATML1 by affecting its activity. We have shown that ATML1 promotes expression of a *LGO* transcriptional reporter at early stages of sepal development (*pLGO::3xVENUS-N7*) (Vadde et al., 2024). We hypothesized that ALE2 might reduce ATML1’s activity and thus its ability to promote *LGO* transcription. To test this, we imaged *pLGO::3xVENUS-N7* in a wild-type background and *pLGO::3xVENUS-N7* in an *ale2-1* mutant background (Figure 7A, B). Because the amount of *LGO* expression changes rapidly in these early developmental stages, we carefully staged buds according to how close the adaxial and abaxial sepals were to enclosing the floral meristem by calculating the distance in pixels between the tips of the abaxial and adaxial sepals. Staging information was incorporated into the graph of the LGO transcriptional reporter expression (Figure 7C). We found that the LGO transcriptional reporter had both reduced and delayed expression in *ale2-1* mutants (Figure 7A–C), similar to what we saw in *atml1-3* mutants (Vadde et al., 2024). Thus, our data are consistent with our hypothesis that ALE2 is necessary for proper ATML1 activity. The fact that overexpressing *ATML1* greatly above endogenous levels in an *ale2-1* background is still able to produce giant cells suggests that ATML1 still retains some function even without ALE2 (Figure 7D). It is possible that ALE2 increases ATML1 activity so that less ATML1 protein is needed to promote LGO transcription at the right time and to the right level (Figure 7D). Hence, we conclude that ALE2 is necessary for proper timing and level of *LGO* transcription at endogenous levels of ATML1.

**Figure 7:**
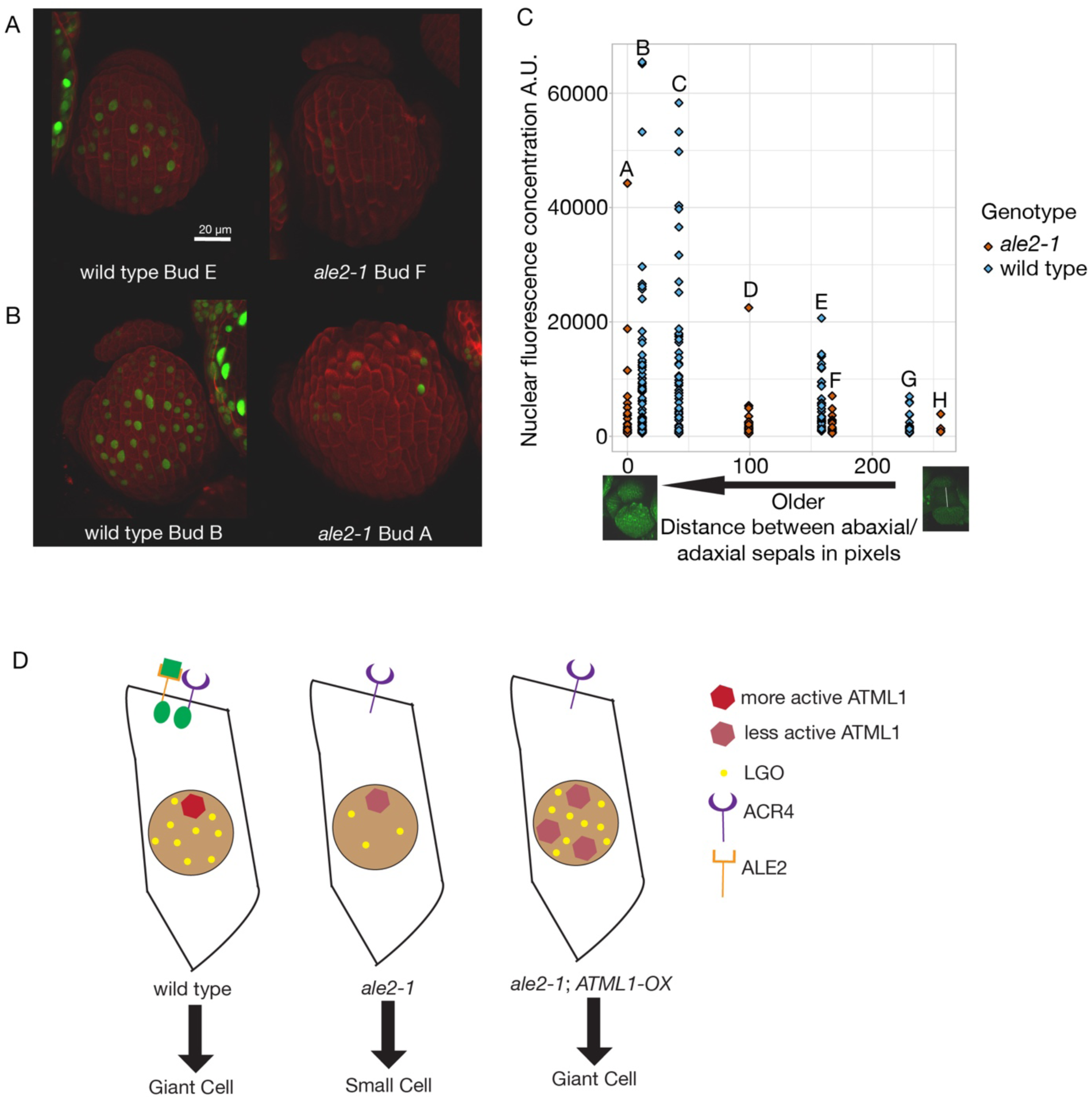
ALE2 is necessary for the timing and strength of *LGO* expression. (A–B) Young buds expressing the *LGO* transcriptional reporter (*pLGO::3xVENUS-N7*) and stained with PI. (A) Wild-type and *ale2-1* mutant bud closely stage-matched. (B) Another wild-type and *ale2-1* mutant bud closely stage-matched. Scale bar is 20 µm. (C) Quantification of the number of nuclei and their fluorescence driven by the LGO transcriptional reporter. Each bud is labeled with a letter. The x axis is distance in pixels between abaxial and adaxial sepals as a method of more closely stage matching the buds. As the sepals grow, they come closer and eventually contact one another (distance 0 pixels), so older buds have less distance between sepals. Buds right next to one another on the graph are very closely stage matched. Buds from (A–B) are included in the graph. (D) Cartoon model of hypothesized effect of ALE2 on ATML1 activity, *LGO* expression, and giant cell formation. In the absence of ALE2, ATML1 activity is reduced and LGO expression decreases. By providing greater than endogenous concentrations of ATML1, LGO expression can increase enough for giant cell specification even in the absence of ALE2. Green square represents a hypothetical ligand for ALE2 and green ovals represent hypothesized cross phosphorylation of ALE2 and ACR4.

## Discussion

Here, we show that ALE2 functions in giant cell specification in Arabidopsis sepals and leaves in addition to its previously known role in epidermal specification. We show that in *ale2-1* mutants there is a loss of enlarged, polyploid cells in addition to a severe loss in giant cell identity based on activity of a giant cell-specific enhancer. We place ALE2 genetically upstream of ATML1 and LGO within the giant cell specification pathway. ALE2 does not affect ATML1 protein concentrations during early sepal development but strongly decreases and delays LGO transcription as determined by a *LGO* transcriptional reporter. Our data is consistent with the model that ALE2 sensitizes cells of the developing sepal epidermis to fluctuations of the transcription factor ATML1 (Figure 7D).

Since *ALE2* encodes a transmembrane receptor-like kinase, signaling is likely to be involved in giant cell patterning. The other receptor-like kinase known to function in giant cell patterning, ACR4, has been shown to both phosphorylate and be phosphorylated by ALE2 *in vitro* (Tanaka et al., 2007). *ale2* and *acr4* mutants also have a similar phenotype in that they have a compromised cuticle and ovule defects (Gifford, Dean, & Ingram, 2003; Tanaka et al., 2007). Thus, ACR4 and ALE2 act in the same pathway, likely to transduce a signal promoting giant cell development. *ACR4* has 4 paralogs in Arabidopsis, whereas *ALE2* does not have any paralogs (Nikonorova et al., 2015). This could explain why the *ale2* mutant phenotype is stronger than the *acr4* mutant phenotype, which is likely to be mitigated by some redundancy of the paralogs. In the root, the ligand of ACR4 has been shown to be CLE40 (Stahl et al., 2009). However, we do not know the identity of its ligand in the sepal. The extracellular ligand binding ALE2 is also unknown and, furthermore, it is unknown from where the signal originates. A signaling ligand could come from other epidermal cells, mesophyll cells, or perhaps even from giant cells themselves in a form of autocrine signaling whereby the signaling ligand comes from the same cell perceiving the signal. Here, we show that the *ALE2* promoter is initially expressed in all epidermal cells but is turned off in giant cells some time after differentiation, suggesting that signaling is no longer required at that point. When *ALE2* expression is forced to remain on in giant cells, giant cell patterning is not perturbed, indicating that turning off signaling is not essential for patterning. Future work is needed to elucidate the identity of the signaling molecule, where it is localized, and which cells perceive it.

Although the details of intercellular signaling remain unknown, we have established the role of ALE2 in the giant cell specification pathway. In the model of giant cell specification, the transcription factor ATML1 fluctuating to high concentrations during G2 of the cell cycle triggers giant cell formation (Meyer et al., 2017). We show that ALE2 fits into this model by being necessary for proper timing and strength of *LGO* expression. *ALE2* functions genetically upstream of *ATML1* but is not necessary for proper ATML1 protein fluctuations. ATML1 protein fluctuates to similarly high peak levels in both *ale2* and wild-type buds. ALE2 must then function upstream of ATML1 by directly or indirectly affecting the ability of the ATML1 protein to function properly. It may do this in a variety of ways. For instance, ALE2 may promote the expression of another gene encoding a protein that helps ATML1 bind to its DNA targets. Or, for example, ALE2 may initiate a phosphorylation cascade ending with the phosphorylation of ATML1 to make it more active. Another possibility is that ALE2 enables the binding of a lipid molecule to ATML1’s lipid-binding START2 domain and that this binding makes ATML1 more active through dimerization (Vadde et al., 2024). In whatever way ALE2 affects ATML1’s ability to function properly, it is clear that ATML1 retains some function even without ALE2. The fact that over-expressing *ATML1* in an *ale2-1* mutant background is still able to produce giant cells indicates that ATML1 is still able to specify giant cells without ALE2 when at much higher than endogenous concentrations. At endogenous ATML1 concentrations, however, it is clear that ALE2 is necessary for ATML1 to specify giant cells. Thus, ALE2 seems to sensitize cells of the developing epidermis to endogenous ATML1 concentration peaks (Figure 7D).

The CDK inhibitor LGO acts downstream of ATML1 and is necessary for high ATML1 concentrations to trigger giant cell formation (Meyer et al., 2017). ATML1 promotes *LGO* transcription (Vadde et al., 2024). In the absence of ATML1, *LGO* transcription is delayed and decreased, while over-expressing *ATML1* greatly increases *LGO* transcription (Vadde et al., 2024). We show that a *LGO* transcriptional reporter is delayed and decreased in an *ale2* mutant background. This finding is consistent with our model of ALE2 enabling proper ATML1 function (Figure 7D). However, we cannot rule out the possibility that ALE2 affects *LGO* transcription independently of ATML1.

We show that *ale2-1;atml1-3* double mutants are homozygous lethal and arrest at the globular stage of embryogenesis. This suggests that ALE2 is upstream of other epidermal specification factors in addition to ATML1. One possible additional epidermal specification downstream of ALE2 is ATML1’s paralog PDF2. ATML1 and PDF2 function redundantly in epidermal specification (Ogawa et al., 2015). Double loss-of-function mutations in *atml1-3* and *pdf2-4* are lethal and cause globular-stage embryonic arrest (Ogawa et al., 2015). If ALE2 is upstream of PDF2 in addition to ATML1 and ALE2 is important for proper PDF2 function in addition to proper ATML1 function, then *ale2-1*;*atml1-3* double mutant lethality is a logical outcome. The fact that *ale2* single mutant is not embryonic lethal is further evidence that ATML1 retains some degree of function in the absence of ALE2.

This study has yielded a more complete picture of how giant cells are patterned in Arabidopsis by showing how ALE2 functions in relation to the transcription factor ATML1 and the CDK inhibitor LGO. Future work should focus on how ALE2 affects ATML1’s ability to function, as well as identifying the ligand that binds ALE2.

## Materials and Methods

### Plant growth

Plants were grown in Percival plant growth chambers set to 60% humidity, 22 °C temperature, and 24-hour light provided by Philips 800 Series 32 Watt fluorescent bulbs (f32t8/tl841) (∼100 μmol m^−2^ s^−1^) after 4°C stratification. Plants were grown in LM111 soil.

### Mutant screening and map based cloning

M2 L*er* seeds mutagenized with ethyl methanesulfonate (EMS) were purchased from Lehle Seeds and screened for loss of giant cells in the sepal under a Zeiss Stemi 2000 stereomicroscope. The *ale2-3* mutation (internal reference number E43-2) was identified through traditional map based cloning. The loss-of-giant-cells phenotype in *ale2-3* was found to co-segregate with the *erecta* phenotype from the L*er* background. Based on previous findings that many loss-of-giant-cell mutants disrupt genes in the epidermal specification pathway, we looked for epidermal specification genes closest to *erecta* on chromosome 2 (Chr2:11208080..11214662) and found the *ALE2* gene (Chr2:8755852..8760298). We PCR amplified and sequenced the *ALE2* from the *ale2-3* (E43-2) mutant plant and identified a G to A mutation at position 849 of the CDS resulting in a premature stop codon in place of W283. We confirmed that this mutation was responsible for the loss-of-giant-cells phenotype observed because the *ale2-1* canonical allele also exhibits the loss-of-giant-cells phenotype and because the loss-of-giant-cells phenotype in *ale2-1* is rescued by expressing a wild-type *ALE2* transgene.

### Transgenes

To create *pALE2::3xVenus-N7* (pLL28), the *ALE2* promoter was PCR amplified with oLL25 (caccAAG ATT CAT AAG GGT AGT TTG TAT CTC CG) and oLL26 (CTC TTA TCT CAA ATC AAA AAC CAC CAT TAG T) and cloned into pENTER D TOPO to create pLL19 (*ALE2* promoter entry clone). To create the pAR336 *GW::3xVenus-N7* destination gateway binary vector conferring Basta resistance in plants, we PCR amplified *3xVenus-N7* with oAR646 (caccggtaccaaaATGGTGAGCAAGGGCGAGGAG) and oAR647 (CTCTAGaTTACTCTTCTTCTTGATC) and cloned into pENTR D TOPO to create pAR334. Then *3xVenus-N7* was excised from pAR334 with KpnI and XbaI and cloned into pSL11 (Hong et al., 2023) also cut with KpnI and XbaI to create *pAR335 GW::3xVenus-N7* in cloning plasmid BJ36. Next, *GW::3XVenus-N7* was cut from pAR335 with NotI and cloned into pMLBart cut with NotI to create the pAR336 *GW::3xVenus-N7* destination gateway binary vector. Finally an LR reaction was used to recombine the pLL19 ALE2 promoter into the pAR336 binary vector to create pLL28 *pALE2::3xVenus-N7*. Col-0 plants were transformed through *Agrobacterium-*mediated floral dipping and T1 plants were selected with Basta.

To express ALE2 specifically in giant cells, we fused the 1 kb giant cell enhancer (Hong et al., 2023; Roeder et al., 2012) upstream of ALE2 (Giant cell enhancer::ALE2; pAR397). The 3.4 Kb genomic *ALE2* gene was amplified using oAR743 (caccaaaATGCGGAACTTTGCGATGCTTCTG) and oAR744 (ATCAGAGCCAGTCACCATTGCCGGA) and cloned into a pCR8/GW/TOPO vector to create pAR396. An LR reaction was performed to recombine pAR396 into the destination vector pAR201 (Roeder et al., 2012) containing the giant cell promoter to generate pAR397. The transgene was transformed into *ale2-1*/+ plants using *Agrobacterium-*mediated floral dipping (Clough & Bent, 1998). Transgenic T1 plants were selected for hygromycin resistance. The T2 generation was genotyped to identify *ale2-1* homozygous and wild-type plants.

To create *35S::ALE2* (pAR398), an LR reaction was performed recombining the pAR396 *ALE2* genomic DNA into the destination vector pB7WG2 containing the *35S* promoter to generate pAR398 *35S::ALE2*, which was then transformed into *ale2-1*/+ plants with *Agrobacterium-* mediated floral dipping (Clough & Bent, 1998). Transgenic plants were selected for Basta resistance. The T2 generation was genotyped to identify *ale2-1* homozygous and wild-type plants.

To generate *ale2-1* plants with the *pML1::H2B-mGFP* epidermal nuclei marker, pAR180 was incorporated into *ale2-1*/+ plants using *Agrobacterium-*mediated transformation (Clough and Bent, 1998) and transgenic plants were selected for Basta resistance.

### Mutant genotyping

The new *ale2-3* (E43-2) mutant phenotype is recessive and can be genotyped by PCR amplification using primers oAR728 (TTGGTTGGAGCTATTTCTATAATCGTGCCATG) and oAR729 (CTATAGAAGCGTTAAGAAGTCAGAGGGTG) followed by a 4 hour restriction enzyme digestion with NcoI.

*ale2-1* plants lack giant cells as the result of a frameshift mutation due to a single base insertion. The mutant phenotype is recessive and can be genotyped by PCR amplification using oAR522 (CAGTTAAGACCTTCACACTCTCCGA) and oAR523 (CATCAAAGTTGTAGGTTCCAGCCA) followed by a 4 hour restriction enzyme digest with XhoI.

*lgo-2* (SALK_039905) plants lack giant cells. The *lgo-2* mutation is the result of a T-DNA insertion and is a recessive mutation. The *lgo-2* mutation can be genotyped using oAR284 (CTTCTCAACCTCTCACTTCTCCAA), oAR285 (CCGAACACCAACAGATAATT), and JMLB2 (TTGGGTGATGGTTCACGTAGTGGG) (Roeder et al., 2010).

*acr4-2* plants have a reduction in giant cells. The *acr4-2* mutation can be genotyped by PCR amplification using oAR232 (TAGTCACTCTGTGGAATGTCTC), oAR233 (GCACCTACAATTCCTCAATCTG), and SLB1 (GCCTTTTCAGAAATGGATAAATAGCCT).

*dek1-4* plants do not form giant cells. The *dek1–4* mutant phenotype is recessive and can be genotyped by PCR amplification using oAR337 (TCCACAGGTAGTTTCTCTTGC) and oAR338 (ACCTTCTCTAATGTTGCGTG), and subsequent sequencing of the product using oAR338 (ACCTTCTCTAATGTTGCGTG).

*atml1-3* (SALK_033408) plants have a severe reduction in giant cells. The *atml1-3* mutation is the result of a T-DNA insertion in the homeodomain. The mutation is semi-dominant and can be genotyped by PCR amplification using oAR272 (CAGGCAGAAGAAAATCGAGAT), oAR273 (GAAACCAGTGTGGCTATTGTT) and JMLB2 (TTGGGTGATGGTTCACGTAGTGGG).

pHM44 (*pATML1::mCitrine-ATML1)* is an mCitrine reporter fused to ATML1 under the *ATML1* promoter. The presence of the transgene can be detected by PCR amplification using oAR272 (CAGGCAGAAGAAAATCGAGAT) and oAR273 (GAAACCAGTGTGGCTATTGTT), which yields a 1641 bp band for the transgene and a 1215 bp band for the wild type genomic copy of ATML1.

*ML1::LGO* is a *LGO* overexpression line, under the *ML1* promoter, which produces ectopic giant cells. The overexpression transgene can be PCR amplified using oAR204 (CTTCCCTTTCTCCTAAGTTCCT) and oAR295 (GATTATCTCACAAGTCGACAC).

*PDF1::FLAG-ATML1 (GIL90-5)* is an ATML1 overexpression line under the PDF1 promoter (San-Bento et al., 2014). Sepals exhibit an upregulation of giant cells. The overexpression gene can be PCR genotyped by amplifying using oAR273 (GAAACCAGTGTGGCTATTGTT) and oHM56 (CATATGGGAGACAGCTTTCTCATACGCG).

The pAR397 construct is a construct expressing genomic *ALE2* under the giant cell promoter in pAR201. The pAR398 construct is a construct expressing *ALE2* under the *35S* promoter in pB7wG2. The *ale2-1* background can be genotyped by PCR amplification with oAR522 (CAGTTAAGACCTTCACACTCTCCGA) and oAR759 (CCTAACCATGCAGCTCAAGAGT), which is specific to the genome, not the transgene, followed by a 4 hour restriction enzyme digestion with XhoI.

### Genetic crosses

To generate *ale2-1* double mutant lines, we crossed pollen from *ale2-1* homozygous plants to giant cell mutants *lgo-2*, *acr4-2*, *dek1-4, atml1-3*. Homozygous mutant plants were found in the F2 generation by genotyping. To generate *ale2-1* mutant lines in the *ATML1-OX* (*pPDF1::FLAG-ATML1*) and *LGO-OX* (*ML1::LGO*) backgrounds, *ale2-1* homozygous plants were crossed to plants homozygous for the *pPDF1::FLAG-ATML1* transgene or *ML1::LGO* transgene. For each transgene cross, in the F2 generation, plants both homozygous for *ale2-1* and having at least one overexpression transgene copy were identified through genotyping.

To assess whether *ale2-1* mutant sepals retained giant cell identity, *ale2-1* homozygous plants were crossed to plants expressing the giant and small cell markers, pAR111 and CS70134 (Hong et al., 2023; Roeder et al., 2012). Plants homozygous for these lines were segregated in the F3 generation.

To assess ATML1 fluctuations in an *ale2-1* background, *ale2-1* homozygous mutant plants were crossed to plants expressing pHM44 (*pATML1::mCitrine-ATML1*) (Meyer et al., 2017). Plants homozygous for the *ale2-1* and *atml1-3* alleles with pHM44 were segregated in the F3 generation.

### Flow cytometry

Flow cytometry was performed as described in Roeder et al., 2010 using an Accuri C6 flow cytometer. Approximately 50 stage 12 sepals were dissected from wild type Col-0 and *ale2-1* mutant plants expressing the pAR180 epidermal nuclei marker (*pML1::H2B-mGFP*). Nuclei were additionally stained with propidium iodide and gated as described previously (Roeder et al., 2010) to separate GFP positive epidermal nuclei from GFP-negative nuclei of the inner sepal.

### Microscopy

Images of floral morphology were taken on a Zeiss Stemi 2000-C microscope using a Canon PowerShot A640 digital camera.

Scanning electron microscopy was performed according to Roeder et al., 2010. Stage 14 flowers were fixed (3.7% formaldehyde, 5% acetic acid, 50% ethanol) and dehydrated through an ethanol series, followed by critical point drying and sepal dissections. Sepals were sputter-coated with platinum palladium and imaged using a Tescan Mira3 FESEM scanning electron microscope.

Confocal imaging was performed on a Zeiss 710 laser scanning confocal microscope. For imaging the giant and small cell enhancer fluorescent reporters, sepals at approximately stage 6 and stage 9, representing early and mid-stage sepals respectively, were stained using propidium iodide (PI) and imaged using a 20× water immersion lens. Stage 12 sepals representing the late stage were stained with PI and imaged under a 10× objective. The small cell marker was excited with a 488 nm laser and emission was collected at 493-518 nm. The giant cell enhancer was excited with a 514 nm laser and 519-568 nm emission wavelengths were collected. PI was simultaneously excited with the 514 laser and 603-650 nm emission was collected. Imaging of the *pALE2::GFP* (pLL28) expression pattern was performed using a 514 nm laser for excitation and 519-564 nm emission wavelengths were collected. The cell walls were visualized with PI with the same laser and 599-650 nm emission wavelengths were collected. For cell size analysis, stage 12 sepals were PI stained and imaged under a 10× objective as above. For *pATML1::mCitrine-ATML1* imaging, inflorescences were dissected down to very young buds around the inflorescence meristem and mounted in 2% agarose plates filled with water. Images were taken of stage 3 and 4 buds using a 20× water immersion lens and zoom of 2.0 with an excitation laser of 514 nm (collection range=519-566 nm) and detector gain of 640.

For imaging of *pLGO::3xVENUS-N7* buds, inflorescences were dissected down to very young stage 3 or 4 buds and stained with PI, then mounted in 2% agarose plates filled with water. Images were taken using a 20× water immersion lens and zoom of 2.0 with an excitation laser of 514 nm (collection range=519-541 nm for 3xVenusN7 and 613-661 nm for PI).

### Image Analysis

Cell size analysis between wild-type and *ale2-1* sepals was performed by processing Stage 12 PI-stained sepal images semi-automatically using a MATLAB module to determine cell area (Cunha, Roeder, & Meyerowitz, 2010; Roeder et al., 2010). For quantification of mCitrine fluorescence intensity from the images of *pATML1::mCitrine-ATML1 atml1-3* in wild-type and *ale2-1* backgrounds, MorphoGraphX (Barbier de Reuille et al., 2015; Strauss et al., 2022) was used to detect nuclei and calculate fluorescence concentration. First, *pATML1::mCitrine-ATML1* images were blurred by running the process “Gaussian Blur” with the parameters: x=0.6, y=0.6, z=0.6. Next, we ran the process “Local Max” with the parameters: x radius=0.5, y radius=0.5 z radius=0.5, start label=2, threshold=1,000, value=60,000. The result was dots marking local maximums according to the parameters. Most nuclei had one dot, however some had more than one. The Voxel Edit tool was used to erase extra dots so that each nucleus had one dot identifying a local maximum. The process “Mesh from local max” was run with the parameter radius=1, creating a 3-D ball within each nucleus. The average fluorescence concentration within each ball was calculated by running the “Heat Map” process. For quantification of *pLGO::3xVENUS-N7* images in wild-type and *ale2-1* backgrounds, MorphoGraphX was used to detect nuclei and calculate fluorescent concentrations in the same way as for *pATML1::mCitrine-ATML1* images. The parameters were held constant between images so that the same fluorescence intensity cutoff was used to identity *LGO*-expressing nuclei for all images. For *pLGO::3xVENUS-N7* images, the following parameters were used for each process: “Gaussian Blur” (x=0.5, y=0.5, z=0.5), “Local Max” (x radius=0.5, y radius=0.5 z radius=0.5, start label=2, threshold=500, value=60,000), “Mesh from local max” (radius=1).

### Assessment of embryo lethality

Siliques were opened with a needle to remove ovules. Ovules were placed in Hoyer’s Solution (2.5g gum Arabica, 33.3g chloral hydrate, 1.66g glycerol, 10ml water) for 3 to 6 hours for clearing before counting under DIC optics with a MetaMorph imaging system. Only siliques in which the majority of embryos were at the torpedo or walking stick stages were counted. Embryos at the globular stage were counted as arrested. Some globular stage embryos were at various stages of degradation and were hard to see. Therefore, ovules that looked empty were also counted as arrested. All *atml1-3* and *dek1-4* single mutant ovules counted as arrested appeared empty. Embryos that had initiated cotyledons and progressed past the globular stage were counted as not arrested. The proportion arrested was determined for each silique. A two-sample t-test was performed to compare the arrested proportions of *ale2-1/+;atml1-3* siliques to *atml1-3* siliques with the null hypothesis that the difference between the two was not different from 0.25. In the same way, a two-sample t-test was performed to compare the arrested proportions of *ale2-1/+;dek1-4* siliques to *dek1-4* siliques with the null hypothesis that the difference between the two was not different from 0.25.

## Acknowledgements

We thank Clint Ko, Lila Luna, Ariel Johnson, Xian Qu, and Xihang Wang for technical assistance with the project.

## Funding

This work was funded by NSF IOS-1553030 (AHKR) and a core grant from the Max Planck Society (NR, PFJ), and NSF DBI-232051 (AHKR and PFJ).

## Notes

### Competing Interest Statement

The authors have declared no competing interest.

## References

Abe, M., Katsumata, H., Komeda, Y., & Takahashi, T. (2003). Regulation of shoot epidermal cell differentiation by a pair of homeodomain proteins in Arabidopsis. Development, 130(4), 635–643. 10.1242/dev.00292

Balkunde, R., Deneer, A., Bechtel, H., Zhang, B., Herberth, S., Pesch, M., Jaegle, B., Fleck, C., & Hulskamp, M. (2020). Identification of the Trichome Patterning Core Network Using Data from Weak ttg1 Alleles to Constrain the Model Space. Cell Rep, 33(11), 108497. 10.1016/j.celrep.2020.108497

Barbier de Reuille, P., Routier-Kierzkowska, A. L., Kierzkowski, D., Bassel, G. W., Schupbach, T., Tauriello, G., Bajpai, N., Strauss, S., Weber, A., Kiss, A., Burian, A., Hofhuis, H., Sapala, A., Lipowczan, M., Heimlicher, M. B., Robinson, S., Bayer, E. M., Basler, K., Koumoutsakos, P., . . . Smith, R. S. (2015). MorphoGraphX: A platform for quantifying morphogenesis in 4D. Elife, 4, 05864. 10.7554/eLife.05864

Clark, F.K., Weissbart, G., Wang, X., Harline, K., Li, C.B., Formosa-Jordan, P., Roeder, A.H.K. (2024). A common pathway controls cell size in the sepal and leaf epidermis leading to a non-random pattern of giant cells. bioRxiv. 10.17605/OSF.IO/RFCWS

Clough, S. J., & Bent, A. F. (1998). Floral dip: a simplified method for Agrobacterium-mediated transformation of Arabidopsis thaliana. Plant J, 16(6), 735–743. 10.1046/j.1365-313x.1998.00343.x

Cunha, A. L., Roeder, A. H., & Meyerowitz, E. M. (2010). Segmenting the sepal and shoot apical meristem of Arabidopsis thaliana. Annu Int Conf IEEE Eng Med Biol Soc, 2010, 5338–5342. 10.1109/IEMBS.2010.5626342

Digiuni, S., Schellmann, S., Geier, F., Greese, B., Pesch, M., Wester, K., Dartan, B., Mach, V., Srinivas, B. P., Timmer, J., Fleck, C., & Hulskamp, M. (2008). A competitive complex formation mechanism underlies trichome patterning on Arabidopsis leaves. Mol Syst Biol, 4, 217. 10.1038/msb.2008.54

Gifford, M. L., Dean, S., & Ingram, G. C. (2003). The Arabidopsis ACR4 gene plays a role in cell layer organisation during ovule integument and sepal margin development. Development, 130(18), 4249–4258. 10.1242/dev.00634

Hong, L., Rusnak, B., Ko, C. S., Xu, S., He, X., Qiu, D., Kang, S. E., Pruneda-Paz, J. L., & Roeder, A. H. K. (2023). Enhancer activation via TCP and HD-ZIP and repression by Dof transcription factors mediate giant cell-specific expression. Plant Cell, 35(6), 2349–2368. 10.1093/plcell/koad054

Hulskamp, M. (2004). Plant trichomes: a model for cell differentiation. Nat Rev Mol Cell Biol, 5(6), 471–480. 10.1038/nrm1404

Johnson, K. L., Degnan, K. A., Ross Walker, J., & Ingram, G. C. (2005). AtDEK1 is essential for specification of embryonic epidermal cell fate. Plant J, 44(1), 114–127. 10.1111/j.1365-313X.2005.02514.x

Jorgensen, P., & Tyers, M. (2004). How cells coordinate growth and division. Curr Biol, 14(23), R1014–1027. 10.1016/j.cub.2004.11.027

Melaragno, J. E., Mehrotra, B., & Coleman, A. W. (1993). Relationship between Endopolyploidy and Cell Size in Epidermal Tissue of Arabidopsis. Plant Cell, 5(11), 1661–1668. 10.1105/tpc.5.11.1661

Meyer, H. M., Teles, J., Formosa-Jordan, P., Refahi, Y., San-Bento, R., Ingram, G., Jonsson, H., Locke, J. C., & Roeder, A. H. (2017). Fluctuations of the transcription factor ATML1 generate the pattern of giant cells in the Arabidopsis sepal. Elife, 6:e19131. 10.7554/eLife.19131

Nikonorova, N., Vu, L. D., Czyzewicz, N., Gevaert, K., & De Smet, I. (2015). A phylogenetic approach to study the origin and evolution of the CRINKLY4 family. Front Plant Sci, 6, 880. 10.3389/fpls.2015.00880

Ogawa, E., Yamada, Y., Sezaki, N., Kosaka, S., Kondo, H., Kamata, N., Abe, M., Komeda, Y., & Takahashi, T. (2015). ATML1 and PDF2 Play a Redundant and Essential Role in Arabidopsis Embryo Development. Plant Cell Physiol, 56(6), 1183–1192. 10.1093/pcp/pcv045

Roeder, A. H., Chickarmane, V., Cunha, A., Obara, B., Manjunath, B. S., & Meyerowitz, E. M. (2010). Variability in the control of cell division underlies sepal epidermal patterning in Arabidopsis thaliana. PLoS Biol, 8(5), e1000367. 10.1371/journal.pbio.1000367

Roeder, A. H., Cunha, A., Ohno, C. K., & Meyerowitz, E. M. (2012). Cell cycle regulates cell type in the Arabidopsis sepal. Development, 139(23), 4416–4427. 10.1242/dev.082925

San-Bento, R., Farcot, E., Galletti, R., Creff, A., & Ingram, G. (2014). Epidermal identity is maintained by cell-cell communication via a universally active feedback loop in Arabidopsis thaliana. Plant J, 77(1), 46–58. 10.1111/tpj.12360

Schwarz, E. M., & Roeder, A. H. (2016). Transcriptomic Effects of the Cell Cycle Regulator LGO in Arabidopsis Sepals. Front Plant Sci, 7, 1744. 10.3389/fpls.2016.01744

Stahl, Y., Wink, R. H., Ingram, G. C., & Simon, R. (2009). A signaling module controlling the stem cell niche in Arabidopsis root meristems. Curr Biol, 19(11), 909–914. 10.1016/j.cub.2009.03.060

Strauss, S., Runions, A., Lane, B., Eschweiler, D., Bajpai, N., Trozzi, N., Routier-Kierzkowska, A. L., Yoshida, S., Rodrigues da Silveira, S., Vijayan, A., Tofanelli, R., Majda, M., Echevin, E., Le Gloanec, C., Bertrand-Rakusova, H., Adibi, M., Schneitz, K., Bassel, G. W., Kierzkowski, D., . . . Smith, R. S. (2022). Using positional information to provide context for biological image analysis with MorphoGraphX 2.0. Elife, 11. 10.7554/eLife.72601

Tanaka, H., Watanabe, M., Sasabe, M., Hiroe, T., Tanaka, T., Tsukaya, H., Ikezaki, M., Machida, C., & Machida, Y. (2007). Novel receptor-like kinase ALE2 controls shoot development by specifying epidermis in Arabidopsis. Development, 134(9), 1643–1652. 10.1242/dev.003533

Trozzi, N., Lane, B., Perruchoud, A., Wang, Y., Hörmayer, L., Ansel, M., Mollier, C., Malivert, A., Clark, F., Reichgelt, T., Roeder, A.H.K., Hamant, O., Boudaoud, A., Kwiatkowska, D., Runions, A., Smith, R.S., Majda, M. (2023). Puzzle cell shape emerges from the interaction of growth with mechanical constraints. bioRxiv. 10.1101/2023.10.01.560343

Vadde, B.V.L., Russell, N.J., Bagde, S.R., Askey, B., Saint-Antoine, M., Brownfield, B., Mughal, S., Apprill, L. E., Khosla, A., Clark, F.K., Schwarz, E.M., Alseekh, S., Fernie, A.R., Singh, A., Schrick K., Fromme, J.C., Skirycz, A., Formosa-Jordan, P., Roeder, A.H.K. (2024). The transcription factor ATML1 maintains giant cell identity by inducing synthesis of its own long-chain fatty acid-containing ligands. bioRxiv. 10.1101/2024.03.14.584694

Watanabe, M., Tanaka, H., Watanabe, D., Machida, C., & Machida, Y. (2004). The ACR4 receptor-like kinase is required for surface formation of epidermis-related tissues in Arabidopsis thaliana. Plant J, 39(3), 298–308. 10.1111/j.1365-313X.2004.02132.x

Wernet, M. F., Mazzoni, E. O., Celik, A., Duncan, D. M., Duncan, I., & Desplan, C. (2006). Stochastic spineless expression creates the retinal mosaic for colour vision. Nature, 440(7081), 174–180. 10.1038/nature04615

