## Supplementary Figure 1 for "The receptor-like kinase ALE2 promotes giant cell formation in the sepal epidermis"

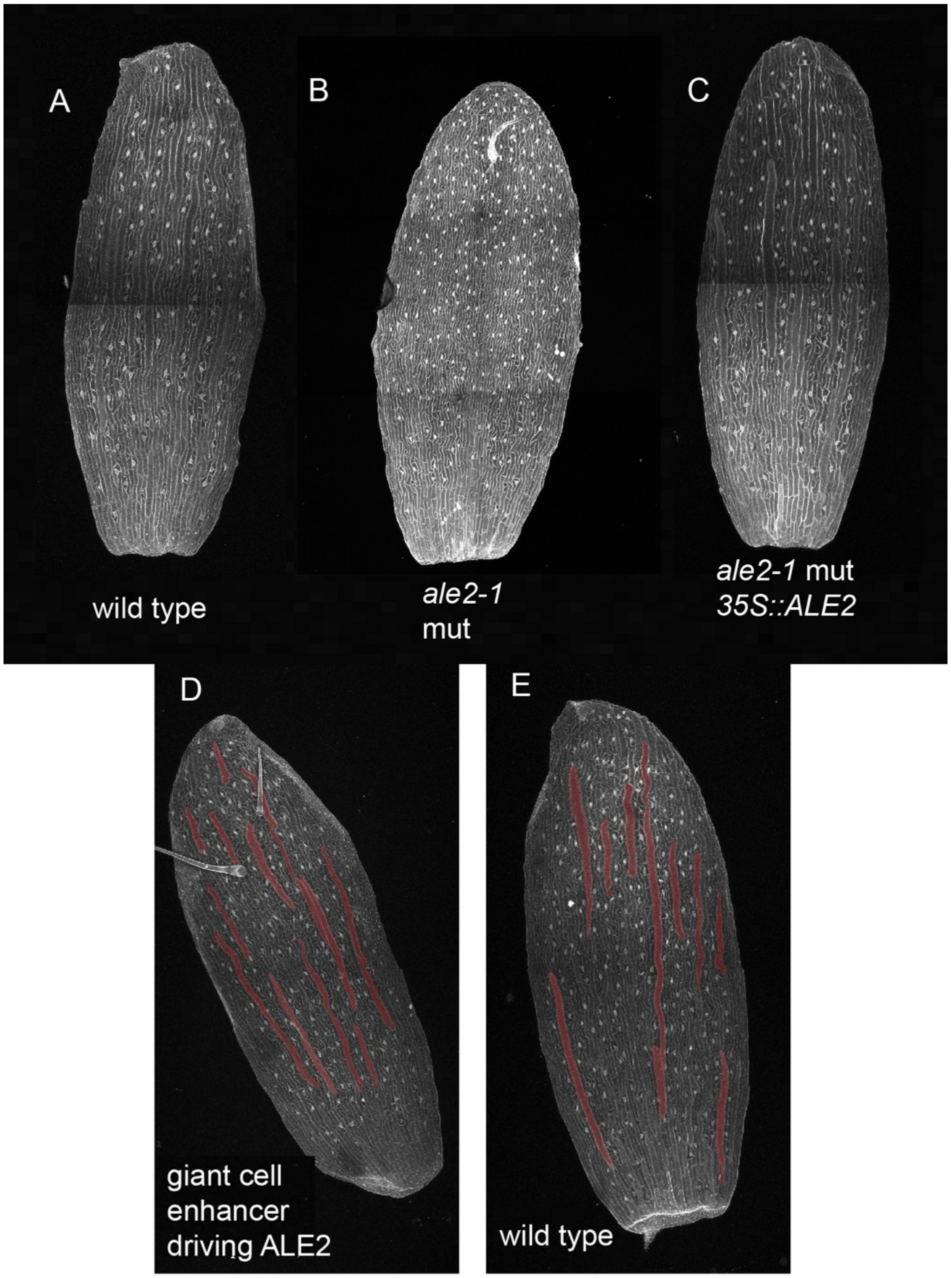


**Supplemental Figure 1: Expression of ALE2 rescues giant cell development in the ale2 mutant**

(A) Wild-type sepal with giant cells. (B) *ale2-1* mutant without giant cells (C) *ale2-1* mutant with *ALE2* being constitutively expressed under the 35S promoter with giant cells apparent. Representative images of (D) wild-type sepal with the giant cell enhancer driving *ALE2* expression. (E) Wild-type sepal without ectopic *ALE2* expression. Giant cells have been false-colored in pink.
